# Automated annotation of rare-cell types from single-cell RNA-sequencing data through synthetic oversampling

**DOI:** 10.1101/2021.01.20.427486

**Authors:** Saptarshi Bej, Anne-Marie Galow, Robert David, Markus Wolfien, Olaf Wolkenhauer

**Author notes:** corrsponding author, website:www.sbi.uni-rostock.de.

## Abstract

The research landscape of single-cell and single-nuclei RNA sequencing is evolving rapidly, and one area that is enabled by this technology, is the detection of rare cells. An automated, unbiased and accurate annotation of rare subpopulations is challenging. Once rare cells are identified in one dataset, it will usually be necessary to generate other datasets to enrich the analysis (e.g., with samples from other tissues). From a machine learning perspective, the challenge arises from the fact that rare cell subpopulations constitute an imbalanced classification problem.

We here introduce a Machine Learning (ML)-based oversampling method that uses gene expression counts of already identified rare cells as an input to generate synthetic cells to then identify similar (rare) cells in other publicly available experiments. We utilize single-cell synthetic oversampling (sc-SynO), which is based on the Localized Random Affine Shadowsampling (LoRAS) algorithm. The algorithm corrects for the overall imbalance ratio of the minority and majority class.

We demonstrate the effectiveness of the method for two independent use cases, each consisting of two published datasets. The first use case identifies cardiac glial cells in snRNA-Seq data (17 nuclei out of 8,635). This use case was designed to take a larger imbalance ratio (∼1 to 500) into account and only uses single-nuclei data. The second use case was designed to jointly use snRNA-Seq data and scRNA-Seq on a lower imbalance ratio (∼1 to 26) for the training step to likewise investigate the potential of the algorithm to consider both single cell capture procedures and the impact of “less” rare-cell types. For validation purposes, all datasets have also been analyzed in a traditional manner using common data analysis approaches, such as the Seurat3 workflow.

Our algorithm identifies rare-cell populations with a high accuracy and low false positive detection rate. A striking benefit of our algorithm is that it can be readily implemented in other and existing workflows. The code basis is publicly available at FairdomHub (https://fairdomhub.org/assays/1368) and can easily be transferred to train other customized approaches.

## 1 Introduction

Single-cell RNA-sequencing (scRNA-Seq), as well as single-nuclei RNA-sequencing (snRNA-Seq), open up a transcriptome-wide gene expression measurement at single-cell level, enabling cell type cluster identification, the arrangement of populations of cells according to novel hierarchies, and the identification of cells transitioning between individual states [1]. This facilitates the investigation of underlying structures in tissue, organism development, and diseases, as well as the identification of unique subpopulations in cell populations that were so far perceived as homogeneous.

### 1.1 Using single-cell technology for the identification of rare cells

Classifying cells into cell types or states is essential for many biological analyses [2]. For example, investigating gene expression changes within a cell type or cell subpopulation can be of high interest across different biological conditions, time-points, or in patient samples. To be able to compare these different cell types, reliable reference systems, especially in sparse-cell states are necessary. However, the lack of markers for rare-cell types motivates the use of unsupervised clustering approaches. Method development for such unsupervised clustering of cells has already reached a certain level of maturity [3, 4, 5]. Furthermore, many studies are interested in specialized cells (e.g., cancer cells, cardiac pacemaker cells) with an occurrence of less than 1 in 1000. The identification of such clusters, solely based on unsupervised clustering of a single dataset, remains very challenging [6]. For this reason, almost every cell clustering characterization approach is driven by manual cluster annotation, which is time consuming and involves a bias of the annotating domain expert, thus limiting the reproducibility of results. One possible solution requires a so-called cell atlas, as a curated reference system that systematically captures cell types and states, either tissue specific or across different tissues [7].

Here, we show how the limitation of identifying already annotated rare-cell types in newly generated scRNA-Seq data can be overcome, by using an synthetic oversampling approach (sc-SynO). sc-SynO is able to automatically identify rare-cell types in an unbiased and precise manner in novel data.

### 1.2 Using machine learning algorithms to generate cell types *in silico*

Machine learning (ML) algorithms are widely used to deal with classification problems and, thus, are used here to automate the annotation of rare-cell types from single-cell or nuclei RNA-Seq data. However, the scarce number of these cells within samples (less than 1 out of 1000 cells) often results in highly imbalanced data. An imbalanced dataset is a type of dataset where one or more classes have a significantly less number of samples compared to other classes (e.g., sinus node cells in the heart). A class having such a low number (minority class) is difficult to detect for unsupervised clustering approaches or classification algorithms in general [6]. The reason behind this is the inability of ML algorithms to perceive or learn underlying patterns from the minority class due to the scarcity of samples and thereby failure of these algorithms to classify them properly [8].

To overcome the problem of imbalance, oversampling techniques have been an area of research in the field of ML for more than a decade. Among several approaches proposed to deal with such issues is the approach of synthetic oversampling [9]. The philosophy of generating synthetic samples is to impute minority class instances, here cell types, in an attempt to enhance the capacity of an ML algorithm to learn. The idea of oversampling is thus commonly used to re-balance the classes [8].

In our study, we compared and benchmarked the ML-based annotation of rare cells with no oversampling and the most commonly used oversampling algorithm **SMOTE** [10], as well as our own sc-SynO approached based on the recently described **LoRAS** algorithm [11]. We provide a brief description of all the oversampling algorithms we have used to pre-process our data as follows:

#### SMOTE

Creates minority class instances by an underlying mathematical assumption. It assumes, that for two close enough minority instances, a convex combination of those two minority instances can be considered as a synthetic minority instance. Generating oversamples with SMOTE has been used to enrich the minority class enabling improvement of ML algorithm performances in case of imbalanced datasets [10].

- **Borderline-SMOTE:** Considers the majority class distribution along with the minority class to look for ‘Borderline’ samples [12].
- **SVM-SMOTE:** Considers the majority class distribution along with the minority class to look for ‘Borderline’ samples, which are selected using support vectors of a pre-trained SVM model. [13].
- **ADASYN:** Decides ‘adaptively’ the number of synthetic samples to be generated from each minority class data point depending on their importance in improving the learning capability of a classifier[14].
- **sc-SynO:** Our proposed approach is based on the LoRAS algorithm and integrates the idea of approximating the minority class data manifold with a more comprehensive modelling of the convex data space of the minority class, while generating synthetic minority class samples resulting in more balanced model performance in terms of F1-Score and Balanced accuracy [11].

## 2 Data and Methods

### 2.1 Use case preparation

To evaluate the potential of synthetic oversampling to precisely annotate cell populations in newly generated data, we generated two use cases by utilizing already published single-cell and nuclei RNA-Seq datasets. Normalized read counts were processed with Seurat [15] (any other normalization method is also applicable). These are then used as an input to generate the synthetic samples and train the different ML classifiers. In addition, we tested the influence of using all transcripts for a classification, or only the top 20, 50, or 100 pre-selected ones (basic feature selection function of Seurat 3 was used). This helps us to investigate the influence of further downstream information obtained from standard feature selection workflows that are usually applied during scRNA-Seq analysis.

The first use case identifies cardiac glial cells in snRNA-Seq data (17 nuclei out of 8,635) [16], which were used as a training set. The trained sc-SynO ML-classifier was subsequently applied to independently generated snRNA-Seq data sets of Wolfien *et al*. [17] and Vidal *et al*. [18] to automatically detect the cardiac glial cells (Glial cells). This use case was designed to take a larger imbalance ratio (∼1 to 500) into account and only uses single-nuclei data.

In contrast to this, the second use case was designed to jointly use snRNA-Seq data and scRNA-Seq on a lower imbalance ratio (∼1 to 26) for the training step to likewise investigate the potential of the algorithm to consider both single cell capture procedures and the impact of “less” rare-cell types. In particular, studies of Galow *et al*. [19] (snRNA-Seq), and Linscheid *et al*. [20] (scRNA-Seq) were used to identify prolifertive cardiomyocytes (Prl cardio). For validation purposes, all datasets have also been analyzed in a traditional manner using common data analysis approaches, such as the Seurat3 workflow, as already described elsewhere [17]. Key statistics of the imbalance ratio (Imb. ratio), number of minority samples, cross-validation folds, and oversampling neighbors of the use cases are presented in Table 1.

**Table 1:** Key statistics of the datasets used during this study. The column ‘Oversampling nbd’ shows the number of nearest neighbours considered for each minority class data points to generate synthetic samples

| Dataset | Imb. ratio | minority samples | CV folds | Oversampling nbd |
| --- | --- | --- | --- | --- |
| Glial cells | 506.94 | 17 | 3 | 3 |
| Prl cardio | 26.21 | 625 | 10 | 30 |

### 2.2 Transferring the LoRAS algorithm to single-cell data

#### The LoRAS algorithm as a basis for sc-SynO

The LoRAS algorithm is designed to create a better approximation of the underlying data manifold by a rigorous modelling of the convex data space compared to pre-existing algorithms, like SMOTE and several of its already presented extensions. A brief outline of the sample generation of sc-SynO approach, as well as the resulting benefits, are shown in Figure 1. To generate a synthetic sample, the algorithm first considers the *k*-nearest neighbours of a minority class data point from a two dimensional embedding of the minority class achieved by using t-SNE. When there are enough data points in the minority class this provides the algorithm a better approximation of the local data manifold for the minority class.

**Figure 1:**
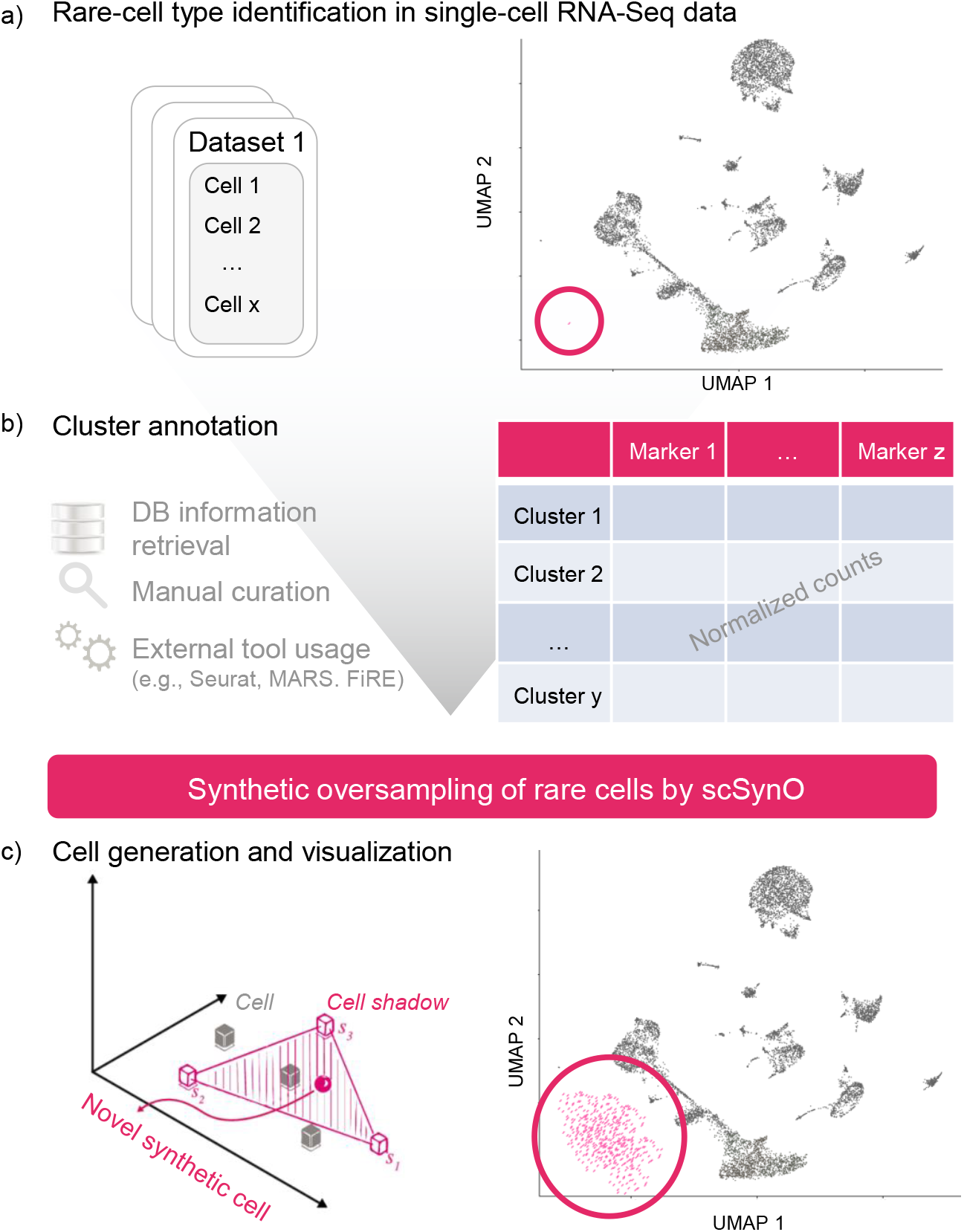
Visualization of the workflow demonstrating a step-by-step explanation for a sc-SynO analysis. **a)** Several or one snRNA-Seq or scRNA-Seq fastq datasets can be used as an input. Here, we identify our cell population of interest and provide raw or normalized read counts of this specific population to sc-SynO for training. **b)** Further information for cluster annotation and processed count data are serving as input for the core algorithm. **c)** Based on the data input, we utilize the LoRAS synthetic oversampling algorithms to generate new cells around the former origin of cells to increase the size of the minority sample. The trained Machine Learning classifier is validated on the trained, pre-annotated dataset to evaluate the performance metrics of the actual model. The model is now ready to identify the learned rare-cell type in novel data.

Once the *k*-nearest neighbours are decided for a minority class data point *p* and thereby the neighbourhood of *p* is identified, Gaussian noise is added to all the data points in the neighbourhood of *p*. The pseudo data points generated by the Gaussian noise are called shadowsamples. A random convex combination of multiple shadowsamples is used to create a Synthetic LoRAS sample. A mathematical explanation of the algorithm asserts that, using convex combination of multiple shadowsamples in LoRAS, can produce a better estimate of the local mean considering the synthetic samples generated in a neighbourhood are random variables [11].

#### ML model description

For our benchmark study, we chose the k-nearest neighbours (knn) and logistic regression model (lr) as our ML classifiers. The reason behind choosing knn is that this model is known to perform well for imbalanced datasets, especially while using oversampling algorithms [21]. We also used the lr model because we observed that the effectiveness of the model in other benchmarking studies using different imbalanced datasets is performing well jointly with the LoRAS oversampling algorithm [11]. The knn model was used with *k* = 30 parameter value. After oversampling there are almost equal data points in the majority and the minority class. For the knn classifier model, we choose a *k* value of thirty to ensure that the classifier’s decision is made on a statistically significant number of samples. The lr model was used with default parameter settings.

Given the proliferative cardiomyocytes dataset, for every ML model, we use a 5 *×* 10-fold stratified cross-validation framework to judge model performances. For the cardiac glial cell dataset, due to the extremely small minority class of only 17 cells, we used a 5*×* 3-fold stratified cross-validation. First, we shuffle the dataset randomly. We divide the dataset into *k*-folds depending on the dataset as described above. The folds are kept distinct maintaining approximately the same imbalance ratio in each fold. After we train and test our models using stratified cross validation, we identify an appropriate model for a given dataset based on the

F1-score and balanced accuracy. The selected model is then trained over the whole dataset and is then used to detect rare cells in two corresponding validation datasets.

#### Oversampling procedure

For our comparative study, we apply the already presented oversampling algorithms. Although there are several other oversampling strategies, convex combination based oversampling can work particularly well when there are too few data points in the minority class due to a lesser chance of overfitting. For every test fold, we oversample only on the training fold, so that the test fold is completely unseen to the classifiers. We specify the most important parameter values of the oversampling model to ensure full reproducibility of our models. For the proliferative cardiomyocytes dataset having 625 minority class samples, we choose 30 of the nearest neighbours of a minority class sample, as the oversampling neighbourhood of all algorithms. sc-SynO has some additional parameters, such as *N*_aff_, *L*_σ_, and *N*_gen_, enabling a better approximation of the minority class data manifold. For the Glial cell dataset, with only 17 minority class samples, we use three of the nearest neighbours of a minority class sample, as well as the oversampling neighbourhood of all algorithms. The *num_afcomb* parameter is chosen to be 23 and 100 for the two cases studies of the proliferative cardiomyocytes dataset with 23 and 100 prioritized marker-genes respectively. For the Glial cell dataset, *num_afcomb* is chosen to be 50 in both case studies. For detailed parameter values please see the code published on FairdomHub (https://fairdomhub.org/assays/1368).

Choosing proper performance metrics are also often a challenge for imbalanced datasets. The usual performance measures such as accuracy or area under the curve (AUC) of receiver operating characteristic (ROC) might be unreliable in this scenario [22]. In our studies, we used two performance measures, the F1-Score (Harmonic mean of precision and recall) and the Balanced accuracy. A lower F1-Score, in our case studies, implies many false negatives and a lower Balanced accuracy implies a lower rate of detection of the minority class. These two measures together can provide a fair understanding of a classifier performance on the datasets we address.

#### Model implementation, execution, and distribution

To allow for an enhanced reusability and transparency of our analysis we provide jupyter notebooks, which can be easily utilized to rerun our analyses or adapt our proposed algorithm to other sc/snRNA-Seq experiments. We used python version 3.7.4. The initial code basis of LoRAS is described in Bej *et al*. [11] and can be accessed at Github (https://github.com/sbi-rostock/LoRAS). To ensure a broad and versatile use of the proposed algorithm, we performed our benchmarking study on a basic personal computer (Processor: Intel(R) Core(TM) i7-8550U CPU @ 1.80GHz, 4 Core(s), 8 Logical Processor(s), 16 GB RAM), which indicates that the actual runtimes on more powerful computers can be shortened significantly.

## 3 Results

Our studies confirm that the pre-selection of features (marker genes) is an important pre-processing. Pre-selection of features not only results in faster models but also produces more reliable results. In Table 2 we show the comparison of runtimes for several pre-selection scenarios using a knn model. We observe a much higher runtime without pre-selection of features. Moreover, the performance of the predictive model on both validation datasets is highly unreliable as in validation dataset VD1 and VD2 there are only 5 and 3 cardiac glial cells respectively as per expert annotations. In contrast, pre-selection of features yields much more accurate results [18, 16]. Without pre-selection the predictive model uses a large number of features leading to an overfitted model. For this reason, we developed our workflow based on pre-selected features obtained from automated feature selection methods in our in-depth comparisons.

**Table 2:** Table showing comparisons among several feature pre-selection scenarios in terms of runtime and efficiency in detection of glial cells for two different validation datasets (VD 1 & 2)

| Dataset | Pre-selection | Pre-processing | Data generation time | Train-test time | Detected/Actual cells |
| --- | --- | --- | --- | --- | --- |
| VD1 | All features | None | - | 11min 56s | 0/5 |
| VD2 | All features | None | - | 2min 19s | 0/3 |
| VD1 | All features | sc-SynO | 4min 3s | 28min 24s | 2423/5 |
| VD2 | All features | sc-SynO | 4min 19s | 5min 8s | 299/3 |
| VD1 | 50 features | sc-SynO | 1.23 s | 1.94 s | 5/5 |
| VD2 | 50 features | sc-SynO | 1.12 s | 484 ms | 3/3 |
| VD1 | 20 features | sc-SynO | 1.12 s | 679 ms | 6/5 |
| VD2 | 20 features | sc-SynO | 1.11 s | 236 ms | 3/3 |

The results for both model training use cases including pre-specified cellular markers and 5-fold stratified cross validation are presented in Table 3.

**Table 3:** Table showing F1-Scores/Balanced accuracies for several oversampling models for our two ML classifiers (lr and knn) and for several numbers of pre-selected features (Marker genes).

| Dataset | ML | Features | Baseline | SMOTE | Bl-1 | Bl-2 | SVM | ADASYN | sc-SynO |
| --- | --- | --- | --- | --- | --- | --- | --- | --- | --- |
| Glial cells | lr | 20 | .924/.965 | .927/.971 | .920/.969 | .909/.983 | .930/.968 | .924/.971 | .921/.971 |
| Glial cells | knn | 20 | .940/.952 | .949/.999 | .923/.98 | .923/.98 | .949/.999 | .949/.999 | .949/.999 |
| Glial cells | lr | 50 | .929/.956 | .957/.981 | .966/.986 | .963/.981 | .957/.981 | .966/.986 | .957/.981 |
| Glial cells | knn | 50 | .910/.923 | .945/.999 | .945/.980 | .945/.980 | .945/.999 | .945/.999 | .945/.999 |
| Prl cardio | lr | 20 | .800/.867 | .646/.932 | .546/.948 | .455/.941 | .564/.952 | .532/.947 | .705/.937 |
| Prl cardio | knn | 20 | .851/.898 | .731/.967 | .66/.966 | .622/.962 | .698/.966 | .613/.963 | .76/.964 |
| Prl cardio | lr | 100 | .867/.919 | .767/.956 | .760/.959 | .630/.962 | .758/.963 | .682/.958 | .854/.940 |
| Prl cardio | knn | 100 | .859/.893 | .756/.970 | .673/.967 | .612/.962 | .710/.968 | .631/.965 | .698/.964 |

### 3.1 sc-SynO achieves a low false negative rate for the identification of glial cells

#### Training data

For the cardiac glial cell dataset all models produce an F1-Score of more than 90 percent. This means that for this dataset irrespective of the used model the risk of getting false negatives is low. However, for the knn models, we observed that the Balanced accuracy is very high for almost all oversampling models including LoRAS. Since this dataset has a extremely rare minority class, only 17 training samples, here detecting the minority class would be more important, especially when the risk of getting false negatives is low. We thus used the knn model with 20 marker genes and LoRAS oversampling to predict on the validation dataset.

#### Validation

We tested the sc-SynO algorithm, which was trained on snRNA-Seq normalized read count data of two independent snRNA-Seq data sets. We identified four out of five cardiac glial cells (Cluster 32) in the first validation set of Wolfien *et al*. and three out of three cells (Cluster15) for the second validation dataset from Vidal *et al*. (Figure 2A, 2B) [18, 16]. In both data sets no false positive predictions have been observed. Figure 2C shows the average gene expression of particular cardiac glial cell markers that are highly expressed in the identified clusters and weakly in other clusters with a close proximity.

**Figure 2:**
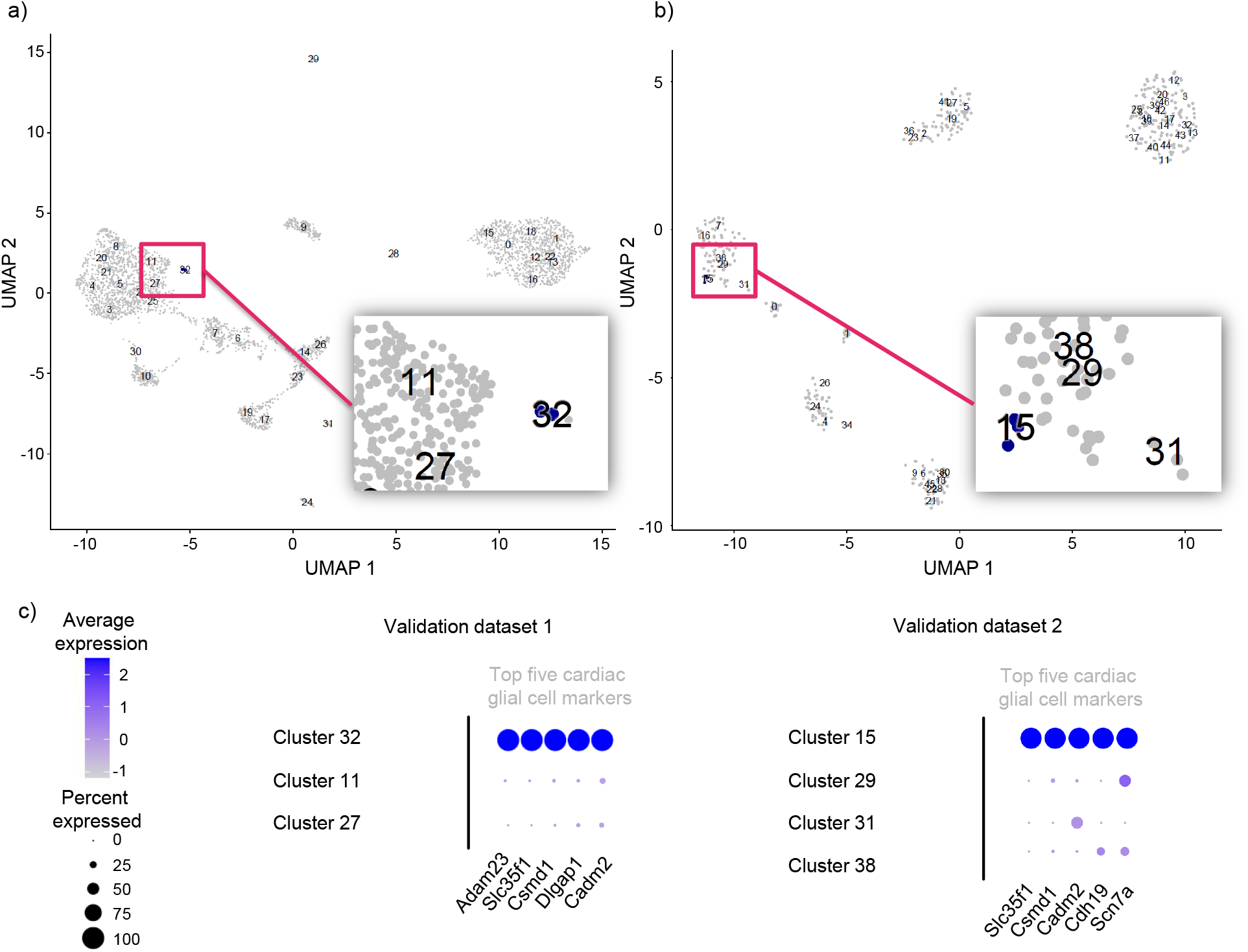
Validation of the sc-SynO model for the first use case of cardiac glial cell annotation. **a)** UMAP representation of the manually clustered Bl6 dataset of Wolfien *et al*. Precicted cells of sc-SynO are highlighted in blue, cells not chosen are grey [17]. **b)** UMAP representation of the manually clustered dataset of Vidal *et al*.. Precicted cells of sc-SynO are highlighted in blue, cells not chosen are grey [18]. **c)** Average expression of the respective top five cardiac glial cell marker genes for both validation sets, including the predicted clusters and those in close proximity.

### 3.2 Oversampling approaches improve the overall identification of proliferative cardiomyocytes

#### Training data

For the proliferative cardiomyocytes dataset, we notice that the performance of the classifiers clearly improves with including more marker genes as features (Figure 3A, 3B). Note that all oversampling models improve on the balanced accuracy compared to the baseline case. However, all of them compromise on the F1-Score. This means, oversampling improves the ability of the classifiers to detect the minority class (the rare-cell type), but renders the classifiers more prone to produce false negatives. Note that for three out of the four case studies, LoRAS compromises the least on the F1-Score compared to the baseline case, while improving on the balanced accuracy. For this dataset, all oversampling models except for LoRAS, produce an F1-Score less than 80 percent, which means they tend to produce more false negatives. Here, the LoRAS algorithm improves on the detection of rare-cell types, while producing less false negatives. Therefore, the lr model with 100 marker genes with LoRAS oversampling was chosen for predictions on the validation dataset.

**Figure 3:**
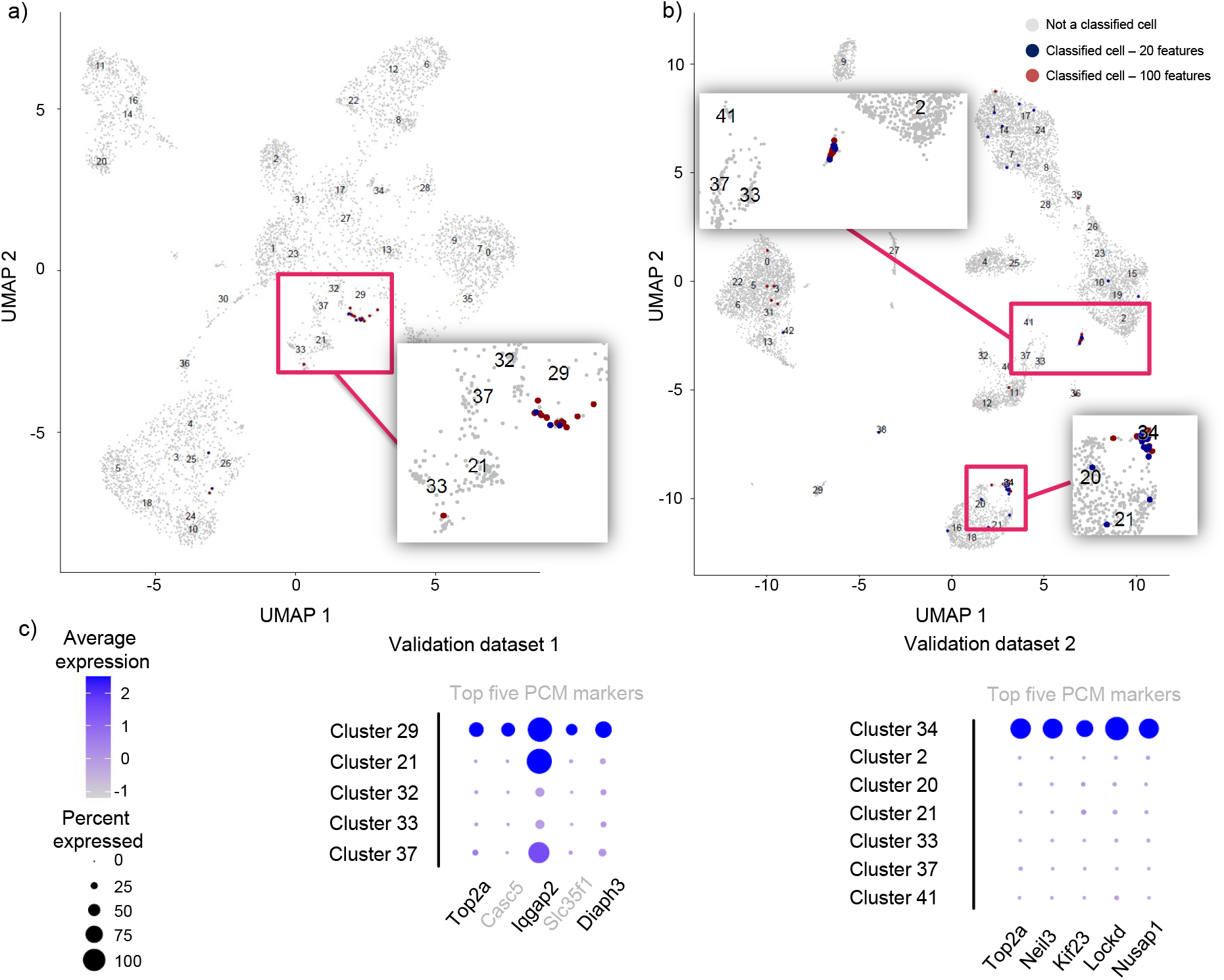
Validation of the sc-SynO model for the second use case of proliferative cardiomyocytes annotation. **a)** UMAP representation of the manually clustered single-nuclei dataset of Linscheid *et al*. Predicted cells of sc-SynO are highlighted in blue (based on top 20 selected features in the training model), red (based on top 100 selected features in the training model) cells not chosen are grey. **b)** UMAP representation of the manually clustered dataset of Vidal *et al*.. Precicted cells of sc-SynO are highlighted in blue (based on top 20 selected features in the training model), red (based on top 100 selected features in the training model) cells not chosen are grey. **c)** Average expression of the respective top five proliferative cardiomyocyte marker genes for both validation sets, including the predicted clusters and those in close proximity.

#### Validation

We applied the sc-SynO algorithm on the two validation datasets for the proliferative cardiomyocytes. We were able to identify 10 out of 11 cells, when using top 20 features. Interestingly, the model using the top 100 genes identifies 48 cells, including 8 common cells with the top 20 model, which may imply that this higher set of transcripts can detect a larger, yet similar, set of cells that are closely related to the cells of investigation. Since the second use case was about a transient cell type, the assigned cells of the model might indicate related cells that have already been or closely to enter the actual state of a proliferative cardiomyocyte. The second validation set assigned 40 cells out of 67 correctly (top 20 features). By using 100 features, the amount of correctly assigned cells increased further up to 340.

## 4 Discussion

Our tool is the first oversampling approach to identify and annotate rare-cell populations from scRNA-Seq and snRNA-Seq data. We already compared and benchmarked the LoRAS algorithm to currently applied other oversampling techniques like SMOTE or Borderline-SMOTE, and present LoRAS as a well-suited algorithm for a broad set of applications in terms of F1-score and balanced accuracy [11]. A principal limitation of SMOTE has been to over-generalize the minority class. This means that the SMOTE algorithm generates many minority class samples, which often are also quite similar to the majority class and divert from the minority class [23]. This results in detection of more minority class samples at the cost of mis-classification of majority class samples as minority class. In other words, the SMOTE algorithm often results in too many false negatives, especially in our use case of single-cell and –nucleus data.

On the other hand, the approach of oversampling using the Borderline samples might lead the algorithms to ignore other important portions of the minority class data space. This causes the oversampled data distribution of the minority class to be too dense in the borderline regions, while too sparse in the other regions. Moreover, oversampling with sc-SynO produces comparatively balanced ML model performances on average, in the sense that, in most cases our algorithm produces less mis-classifications on the majority class with a reasonably small compromise for mis-classifications on the minority class.

Our tool facilitates the identification of very similar cells for smaller sets of feature genes and biologically related cells for larger sets of genes. The initial clustering of the training data plays an essential role, in which we observed that smaller clusters with a distinct border to other clusters are better suited for an analysis in comparison to larger cell populations with transient borders. However, the algorithm still has high accuracies in identifying those cells, but the rate of false positive predictions likewise increases.

In comparison to other current tools, such as cscGAN [24], MARS [25], FiRE [26], and ELSA [27], sc-SynO uses synthetic oversampling of previously, manually curated cell populations to identify such rare cells in novel data. In addition, sc-SynO is easily applicable and only requires a single, well-curated dataset, including only a few cells of interest, to be able to achieve already a high predictive accuracy. Likewise, our algorithm can be used on integrated datasets as well, which commonly represent the underlying biological heterogeneity of the sample in an improved manner [17].

Our approach can be seamlessly incorporated in single-cell and single-nuclei data analysis workflows after the identification and annotation of cell populations on raw or normalized read count data. Once a rare-cell population has been identified and carefully checked by a domain expert, sc-SynO can be used on this manually curated dataset to train the specific cell type. Applying sc-SynO on a novel dataset to identify the same rare-cell type is magnitudes less time-consuming than manually curated data processing and annotation of scRNA-Seq data.

## Availability of data and materials

The computational scripts can be obtained from our FairdomHub instance (https://fairdomhub.org/assays/1368). The single-cell RNA-Seq data utilized during this study is already publicly available at the Single Cell Expression Atlas via ArrayExpress (E-MTAB-7869, E-MTAB-8751, E-MTAB-8848)

## Author contributions

Conceptualization, S.B. and M.W.; methodology, S.B., A.-M.G., and M.W.; formal analysis, S.B., A.-M.G., and M.W.; investigation, S.B., A.-M.G., R.D., M.W. and O.W.; resources, R.D. and O.W.; data curation, S.B. and M.W.; writing the original draft, S.B. and M.W.; writing, review, and editing, S.B., A.-M.G., R.D., M.W. and O.W.; visualization, S.B., M.W., O.W.; supervision, R.D. and O.W.; project administration, R.D. and O.W.; funding acquisition, M.W., O.W., and R.D. All authors have read and agreed to the published version of the manuscript.

## References

[1] D. Lähnemann, J. Köster, E. Szczurek, D. J. McCarthy, S. C. Hicks, M. D. Robinson, C. A. Vallejos, K. R. Campbell, N. Beerenwinkel, A. Mahfouz, et al., “Eleven grand challenges in single-cell data science,” Feb 2020.

[2] J. Lee, D. Hyeon, and D. Hwang, “Single-cell multiomics: technologies and data analysis methods,” Experimental & Molecular Medicine, pp. 1428–1442, Sep 2020.

[3] A. Duò, M. Robinson, and C. Soneson, “A systematic performance evaluation of clustering methods for single-cell rna-seq data [version 2; peer review: 2 approved],” F1000Research, vol. 7, no. 1141, 2018.

[4] S. Freytag, L. Tian, I. Lönnstedt, M. Ng, and M. Bahlo, “Comparison of clustering tools in r for medium-sized 10x genomics single-cell rna-sequencing data [version 2; peer review: 3 approved],” F1000Research, vol. 7, no. 1297, 2018.

[5] V. Y. Kiselev, T. S. Andrews, and M. Hemberg, “Challenges in unsupervised clustering of single-cell rna-seq data,” Nature reviews. Genetics, vol. 20, p. 273—282, May 2019.

[6] A. Jindal, P. Gupta Jayadeva, and D. Sengupta, “Discovery of rare cells from voluminous single cell expression data,” Nature Communications, vol. 9, 12 2018.

[7] F. Zhang, B. Lehallier, N. Schaum, and T. Q. Li, “Single-cell transcriptomics of 20 mouse organs creates a tabula muris,” Nature, vol. 562, pp. 367–372, 10 2018.

[8] B. Santoso, H. Wijayanto, K. A. Notodiputro, and B. Sartono, “Synthetic over sampling methods for handling class imbalanced problems: A review,” IOP Conference Series: Earth and Environmental Science, vol. 58, pp. 012–031, Mar 2017. doi: https://doi.org/10.1088/1755-1315/58/1/012031, ISSN: 1755-1315.

[9] G. Weiss, K. McCarthy, and B. Zabar, “Cost-sensitive learning vs. sampling: Which is best for handling unbalanced classes with unequal error costs?,” in DMIN, pp. 35–41, 01 2007.

[10] N. V. Chawla, K. W. Bowyer, L. O. Hall, and W. P. Kegelmeyer, “Smote: Synthetic minority over-sampling technique,” Journal of Artificial Intelligence Research, vol. 16, pp. 321–335, 2002. doi: https://doi.org/10.1613/jair.953.

[11] S. Bej, N. Davtyan, M. Wolfien, M. Nassar, and O. Wolkenhauer, “Loras: An oversampling approach for imbalanced datasets,” arXiv 1908.08346v4, vol. (accepted for publication in Machine Learning (Springer)), 2020. ArXiv link: https://arxiv.org/pdf/1908.08346.pdf.

[12] H. Han, W.-Y. Wang, and B.-H. Mao, “Borderline-smote: A new over-sampling method in imbalanced data sets learning,” in Advances in Intelligent Computing. ICIC, vol. 3644, pp. 878–887, Springer Berlin Heidelberg, 2005. doi: https://doi.org/10.1007/1153805_91, ISBN: 978-3-540-31902-3.

[13] H. M. Nguyen, E. Cooper, and K. Kamei, “Borderline over-sampling for imbalanced data classification,” Int. J. Knowl. Eng. Soft Data Paradigms, vol. 3, pp. 4–21, 2011.

[14] H. Haibo, B. Yang, E. Garcia, and L. Shutao, “Adasyn: Adaptive synthetic sampling approach for imbalanced learning,” in 2008 IEEE International Joint Conference on Neural Networks, June, 2008. doi: https://doi.org/10.1109/IJCNN.2008.4633969, ISBN: 2161-4393.

[15] A. Butler, P. Hoffman, P. Smibert, E. Papalexi, and R. Satija, “Integrating singlecell transcriptomic data across different conditions, technologies, and species,” Nature biotechnology, vol. 36, 05 2018.

[16] M. Wolfien, A.-M. Galow, P. Müller, M. Bartsch, R. M. Brunner, T. Goldammer, O. Wolkenhauer, A. Hoeflich, and R. David, “Single-nucleus sequencing of an entire mammalian heart: Cell type composition and velocity,” Cells, vol. 9, no. 2, 2020.

[17] M. Wolfien, A.-M. Galow, P. Müller, M. Bartsch, R. M. Brunner, T. Goldammer, O. Wolkenhauer, A. Hoeflich, and R. David, “Single nuclei sequencing of entire mammalian hearts: strain-dependent cell-type composition and velocity,” Cardiovascular Research, vol. 116, pp. 1249–1251, 04 2020.

[18] R. Vidal, J. U. G. Wagner, C. Braeuning, C. Fischer, R. Patrick, L. Tombor, M. Muhly-Reinholz, D. John, M. Kliem, T. Conrad, N. Guimarães-Camboa, R. Harvey, S. Dimmeler, and S. Sauer, “Transcriptional heterogeneity of fibroblasts is a hallmark of the aging heart,” JCI Insight, vol. 4, 11 2019.

[19] A.-M. Galow, M. Wolfien, P. Müller, M. Bartsch, R. Brunner, A. Hoeflich, O. Wolkenhauer, R. David, and T. Goldammer, “Integrative cluster analysis of whole hearts reveals proliferative cardiomyocytes in adult mice,” Cells, vol. 9, p. 1144, 05 2020.

[20] N. Linscheid, S. J. R. J. Logantha, P. C. Poulsen, S. Zhang, M. Schrölkamp, K. L. Egerod, J. J. Thompson, A. Kitmitto, G. Galli, M. J. Humphries, H. Zhang, T. H. Pers, J. V. Olsen, M. Boyett, and A. Lundby, “Quantitative proteomics and single-nucleus transcriptomics of the sinus node elucidates the foundation of cardiac pacemaking,” Nature Communications, vol. 10, no. 1, p. 2889, 2019.

[21] R. Blagus and L. Lusa, “Smote for high-dimensional class-imbalanced data,” BMC Bioinformatics, vol. 14, p. 106, Mar 2013. doi: https://doi.org/10.1186/1471-2105-14-106, ISSN: 1471-2105.

[22] T. Saito and M. Rehmsmeier, “The precision-recall plot is more informative than the roc plot when evaluating binary classifiers on imbalanced datasets,” PLOS ONE, vol. 10, pp. 1–21, 03 2015. doi: https://doi.org/10.1371/journal.pone.0118432.

[23] K. Puntumapon and K. Waiyamai, “A pruning-based approach for searching precise and generalized region for synthetic minority over-sampling,” in Advances in Knowledge Discovery and Data Mining, (Berlin, Heidelberg), pp. 371–382, Springer Berlin Heidelberg, 2012.

[24] M. Marouf, P. Machart, S. Magruder, V. Bansal, C. Kilian, C. Krebs, and S. Bonn, “Realistic in silico generation and augmentation of single cell rna-seq data using generative adversarial neural networks,” Nature Communications volume, vol. 11, p. 166, 08 2018.

[25] M. Brbić, M. Zitnik, S. Wang, A. Pisco, R. Altman, S. Darmanis, and J. Leskovec, “Mars: discovering novel cell types across heterogeneous single-cell experiments,” Nature Methods, pp. 1–7, 10 2020.

[26] A. Jindal, P. Gupta Jayadeva, and D. Sengupta, “Discovery of rare cells from voluminous single cell expression data,” Nature Communications, vol. 9, 12 2018.

[27] L. Wang, F. Catalan, K. Shamardani, H. Babikir, and A. Diaz, “Ensemble learning for classifying single-cell data and projection across reference atlases,” Bioinformatics, vol. 36, pp. 3585–3587, 02 2020.

